# Disrupted myelin homeostasis in grey matter of Alzheimer’s disease

**DOI:** 10.64898/2026.07.08.737146

**Authors:** Assia Tiane, Emily Willems, Lisa Koole, Melissa Schepers, Daniel L. A. van den Hove, Tim Vanmierlo

**Author notes:** These authors contributed equally to this work (Daniel L. A. van den Hove, Tim Vanmierlo).

## Abstract

Alzheimer’s disease (AD) is characterized not only by amyloid-β plaques, tau neurofibrillary tangles and associated neuronal loss, but also by alterations in non-neuronal cell types essential for neuronal support. Oligodendrocytes and their myelin sheaths play a central role in maintaining axonal function, yet detailed molecular profiling of myelin dynamics in the human AD brain remains limited. Although neuroimaging studies increasingly highlight myelin degeneration in white matter as an important contributor to AD pathophysiology, the status of myelin within cortical grey matter is less well understood.

Here, we performed a detailed histopathological characterization of myelin integrity and oligodendrocyte dynamics in both grey and white matter of the middle temporal gyrus (MTG), making use of post-mortem tissue from AD cases (n = 15) and age-, sex-, and *APOE* genotype–matched controls (n = 15). Strikingly, we identified a specific vulnerability of cortical grey matter myelin in AD, whereas white matter myelin appeared relatively preserved. This selective grey matter disruption was accompanied by a seemingly insufficient oligodendrocyte regenerative response, suggesting ongoing attempts at myelin repair, yet featured by a differentiation block. Importantly, the extent of myelin damage and OPC differentiation strongly correlated with proximity to tau pathology, linking cortical demyelination to neuronal and synaptic dysfunction within vulnerable AD regions.

Together, our findings reveal cortical grey matter myelin disruption as a previously underrecognized, highly localized feature of AD pathology. By highlighting the tight intertwining of oligodendrocyte and myelin dynamics with tau-associated neurodegeneration, this work positions cortical myelin pathology as a potential new mechanistic and therapeutic avenue in AD.

## Introduction

Alzheimer’s disease (AD) is classically defined by the accumulation of amyloid-β plaques and neurofibrillary tau tangles. However, the disease is increasingly recognized as a multifaceted disorder involving profound alterations in non-neuronal cell types (1). Among these, oligodendrocytes and their myelin sheaths play a critical yet underappreciated role in maintaining axonal integrity, metabolic support, and efficient neural network function (2, 3).

Over the past decade, neuroimaging studies have consistently reported early widespread white matter abnormalities in individuals with AD, including reduced white matter integrity and altered connectivity (4–6). These findings suggest that myelin disruption is not merely a secondary consequence of neurodegeneration, but may represent an early and contributing factor in disease progression. In parallel, experimental work in mouse models has provided evidence for oligodendrocyte dysfunction and impaired myelin maintenance in AD, including the emergence of disease-associated oligodendrocyte states (7–10). Together, these studies point toward a dynamic process of myelin remodeling and degeneration in the diseased brain.

Despite these advances, detailed molecular and histopathological insights into myelin alterations in the human AD brain remain limited. Most of the current understanding is derived from neuroimaging or experimental model systems, while systematic analyses of myelin pathology in human post-mortem brain tissue remain underexplored (9, 11). As a result, the extent, regional specificity, and cellular underpinnings of myelin alterations across the spectrum of AD neuropathology remain insufficiently characterized. Importantly, existing histopathological myelin studies have rarely been integrated with quantitative neuropathological staging, such as amyloid and tau burden or Braak staging, limiting our ability to position myelin changes within the temporal and spatial progression of the disease.

In this study, we sought to address this gap by systematically examining myelin integrity and oligodendrocyte dynamics in human post-mortem brain tissue. In particular, we focused on delineating differences in myelination between grey and white matter, aiming to identify region-specific patterns of vulnerability. By comparing age-, sex-, and *APOE* genotype–matched control and AD cases, we identified a pronounced susceptibility of grey matter regions to myelin disruption, highlighting that myelin pathology in AD is not confined to classical white matter tracts but also prominently affects intracortical myelinated networks. Together, our findings characterize myelin alterations and define the molecular programs associated with myelin maintenance and disruption in the human AD brain.

## Methods

### Patient cohort

Human post-mortem brain tissue was obtained from the Banner Sun Health Research Institute Brain and Body Donation Program (BSHRI-BBDP; Arizona, USA) and subsequently stored at the Central Biobank MUMC+ (Maastricht, The Netherlands). Middle temporal gyrus (MTG) samples were selected from non-neurological controls (n = 15) and patients with AD (n = 15). Groups were matched for age, sex, and *APOE* genotype. Detailed demographic and clinical characteristics are provided in Table 1.

**Table 1:** Demographic characteristics of the study cohort.

|  | Sex | Age, mean (SD) | APOE | MMSE, mean (SD) | Braak, mean (SD) | PMI, mean (SD) |
| --- | --- | --- | --- | --- | --- | --- |
| **Control** | 8F, 7M | 86 ($\pm 4.8$) | 5 (2/3); 5 (3/3); 5 (3/4) | 27.8 ($\pm 1.5$) | 2.5 ($\pm 1$) | 2.9h ($\pm 0.8$) |
| **AD** | 8F, 7M | 87 ($\pm 7.5$) | 5 (2/3); 5 (3/3); 5 (3/4) | 14.1 ($\pm 9.9$) | 5.1 ($\pm 0.7$) | 3.0h ($\pm 0.7$) |

AD: Alzheimer's disease; F: female; M: male; APOE: Apolipoprotein E genotype; MMSE: mini mental state examination; PMI: post-mortem interval

The BBDP enables rapid post-mortem tissue collection, resulting in a median post-mortem interval of approximately 3 hours (12). In addition to comprehensive ante-mortem neurological and neuropsychological assessments, extensive neuropathological evaluations were performed at BSHRI. Large frozen tissue blocks were stained using Campbell–Switzer, Gallyas, and Thioflavine S methods for the assessment of amyloid plaques, neurofibrillary tangles, and other pathological inclusions (13). These assessments provide a broad range of clinical and pathological data, including mini mental state examination (MMSE) scores, Braak staging, neurofibrillary tangle load, and amyloid plaque burden.

### Immunofluorescence

Frozen brain samples were sectioned into 10 µm-thick cryosections and stored at −20 °C until further use. Sections were used for the assessment of myelin integrity (myelin basic protein [MBP], degraded MBP [dMBP]) and oligodendrocyte phenotyping (2’,3’-cyclic nucleotide 3’-phosphodiesterase [CNPase], Adenomatous Polyposis Coli [CC1]/Oligodendrocyte Transcription Factor 2 [OLIG2], Breast Carcinoma Amplified Sequence 1 [BCAS1], and Phospho-Tau (Ser202, Thr205) [AT8]).

Prior to staining, all sections were air-dried for 30 minutes at room temperature. For dMBP, BCAS1, and AT8 immunostaining, sections were rehydrated in PBS and fixed in ice-cold acetone for 10 minutes. For MBP, CNPase, and CC1/OLIG2, antigen retrieval was performed by incubation in boiling citrate buffer for 30 minutes, followed by two washing steps in 0.1% PBS-T. All sections were subsequently blocked in 10% protein block (Dako) prepared in 0.1% PBS-T for 1 hour, or for 2 hours in the case of antigen-retrieved sections. Primary antibodies were diluted in blocking solution and incubated overnight at 4 °C in a humidified chamber. Sections were subsequently washed three times with PBS and incubated with secondary antibodies diluted in PBS for 1 hour at room temperature. Detailed antibody information is provided in Supplementary Table S1. Following three additional PBS washes, nuclei were counterstained with 4′,6-diamidino-2-phenylindole (DAPI; Invitrogen) for 10 minutes. Sections were then mounted using a fluorescent mounting medium (Invitrogen).

### Imaging and image analysis

Tissue sections were scanned at 20× magnification using the Axioscan Z.1 slide scanner (Zeiss). Whole-slide images were exported and analyzed using QuPath (version 0.5.1). Within each region of interest (ROI; grey and white matter, respectively), three square annotations of 1 mm² were defined for quantitative analysis. Depending on the marker, either the positive area or the percentage of positive cells was assessed within each ROI. The positive area was quantified by applying a pixel classifier and normalizing the classified positive signal to the total annotated area. For cell-based analyses, nuclei were detected using DAPI staining with the cell detection function in QuPath. A single-measurement classifier was subsequently applied to define a threshold for marker-positive cells, enabling the calculation of the percentage of positive cells within each annotation. The mean value of the three annotations was used as a single data point per sample. For the peri-tau BCAS1^+^ analysis, four concentric rings were annotated around each AT8^+^ tangle, with each ring extending 200 µm further from the tangle. The number of BCAS1^+^ cells within each ring was quantified as described above. To determine the number of cells specifically present within each distance interval, the cell count of the preceding inner ring was subtracted from that of the subsequent outer ring. Five peri-tau regions were analyzed per sample, and the resulting values were averaged to obtain a single data point for each sample.

### RNA isolation and transcriptomics

Total RNA was isolated using a standard TRIzol-chloroform extraction protocol (TRIzol, Invitrogen). Library preparation was performed by Novogene using the Novogene NGS RNA Library Prep Set and sequencing was carried out on the NovaSeq X Plus platform.

Raw sequencing reads (FASTQ format) were initially processed using fastp to remove adapter sequences, poly-N reads, and low-quality reads, yielding high-quality clean reads. Clean paired-end reads were aligned to the human reference genome (GRCh38.p14) using HISAT2 (v2.2.1), after building the corresponding genome index. Resulting BAM alignment files were processed in Rstudio (v2026.01) using Rsamtools (v2.18). Gene-level read counts were generated using Rsubread (v2.16.1). Genes with fewer than 10 counts in more than 50% of samples were filtered out, leaving 17,323 genes for downstream analysis. Raw gene counts were normalized using the trimmed mean of M-values (TMM) method using EdgeR (v4.0.16). Differential gene expression (DGE) analysis was performed using DESeq2 (v1.42.1), with age, sex, and RNA integrity number (RIN) included as covariates in the design. Gene ontology (GO) overrepresentation analysis and gene set enrichment analysis (GSEA) were performed using the enrichGO and GSEA functions, respectively, from clusterProfiler (v4.10.1).

To quantify enrichment of a grey matter–specific mature oligodendrocyte transcriptional signature, we performed single-sample gene set enrichment analysis (ssGSEA) using the GSVA package in R. A gene set representing mature oligodendrocytes was derived from a publicly available RNA sequencing dataset (Gene Expression Omnibus (GEO), accession number GSE281006) by identifying genes differentially expressed in grey matter oligodendrocyte populations, compared to white matter oligodendrocytes (14). This gene set was subsequently used as a reference gene set for our bulk RNA sequencing data from grey and white matter control and AD brain samples. ssGSEA was performed with default parameters, as implemented in GSVA, generating sample-wise enrichment scores reflecting the relative activity of the gene signature in each sample. Enrichment scores were compared between AD and control groups within grey and white matter to assess disease- and tissue-specific differences in oligodendrocyte-related transcriptional activity.

### Statistics

Statistical analyses were performed using GraphPad Prism v10 (GraphPad Software Inc.). Differences between groups were assessed using the non-parametric Mann–Whitney test. Correlation analyses were performed using the Spearman rank correlation test. Data are presented as mean ± standard error of the mean (SEM). Statistical significance was defined as p < 0.05 (* p<0.05, **p<0.01), while trends were defined as p ≤ 0.1.

## Results

### Selective myelin alterations in grey matter are associated with disease progression

To assess compact myelin in post-mortem brain tissue from AD patients and controls, we performed immunofluorescent staining for MBP (Figure 1A,B). Quantification of MBP immunoreactivity, expressed as the average MBP-positive area per mm², revealed a significant (*P =* 0.0024) reduction within grey matter regions of AD patients compared to controls. This reduction showed a trend toward association with Braak stage (*P* = 0.0681, r = −0.3563) and MMSE score (*P* = 0.0777, r = 0.3521) (Figure 1C–E), but was not associated with amyloid plaque burden (Supplemental Figure S1). In contrast, no significant differences were detected within white matter regions. However, both control and AD white matter exhibited a relatively low overall MBP-positive area (approximately 50%), suggesting the presence of white matter alterations that are not disease-specific, as they also did not correlate with clinical or pathological markers (Figure 1F–H).

**Figure 1.**
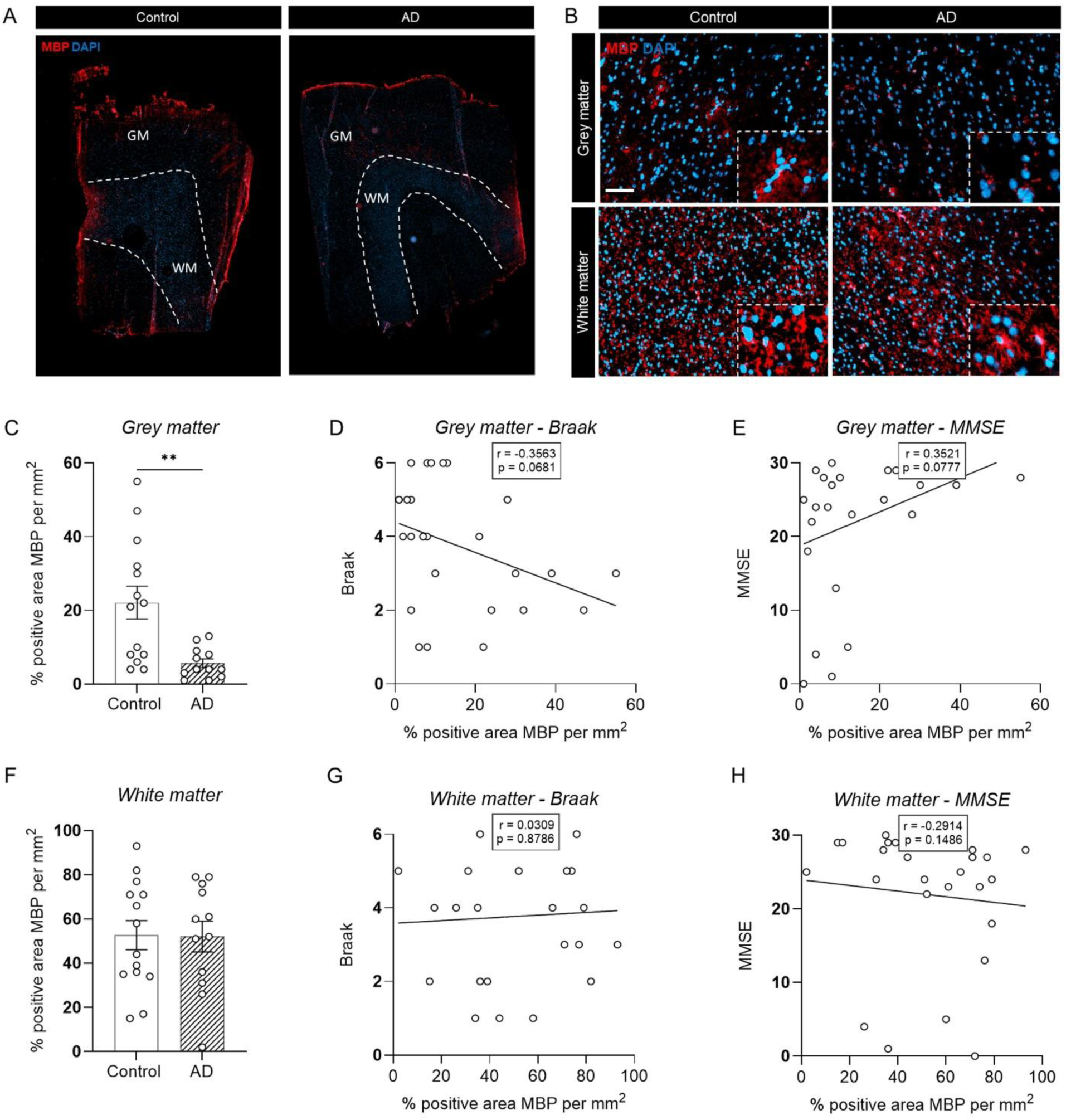
Reduced compact myelin within grey, but not white matter, in AD patients. **(A–B)** Representative myelin basic protein (MBP) staining in brain sections from AD patients and age-matched controls, highlighting grey and white matter regions (scale bar: 50 µm). **(C, F)** Quantification of MBP-positive area (fluorescence area per mm²) revealed a significant reduction of compact myelin in the grey matter of AD patients compared to controls, while no difference was observed in white matter (n = 12–14, Mann–Whitney test, **p < 0.01). **(D–E)** Correlation analyses showed a trend between grey matter MBP area (%) and both Braak stage and mini mental state examination (MMSE) scores (Spearman correlation). **(G–H)** No significant correlations were observed between white matter MBP area (%) and Braak stage or MMSE score (Spearman correlation).

Next, we employed a specific antibody that selectively recognizes myelin degeneration by targeting degraded MBP (dMBP) (Figure 2A,B). This approach enabled quantification of myelin damage in AD and control brain samples. We observed a two-fold increase in the average grey matter dMBP-positive area in AD patients relative to controls (Figure 2C). Notably, this increase in myelin damage was positively associated with Braak stage (*P* = 0.0232, r = 0.4355), showed a trend toward a negative association with MMSE score (*P* = 0.0703, r = −0.3607) (Figure 2D,E), and was positively associated with tangle load (*P* = 0.0330, r = 0.4114), but not with plaque load (Supplementary Figure S1). In contrast, no significant differences nor associations were observed within white matter regions when comparing AD and control samples (Figure 2F–H). Notably, a considerable degree of white matter myelin damage seemed present in both groups.

**Figure 2.**
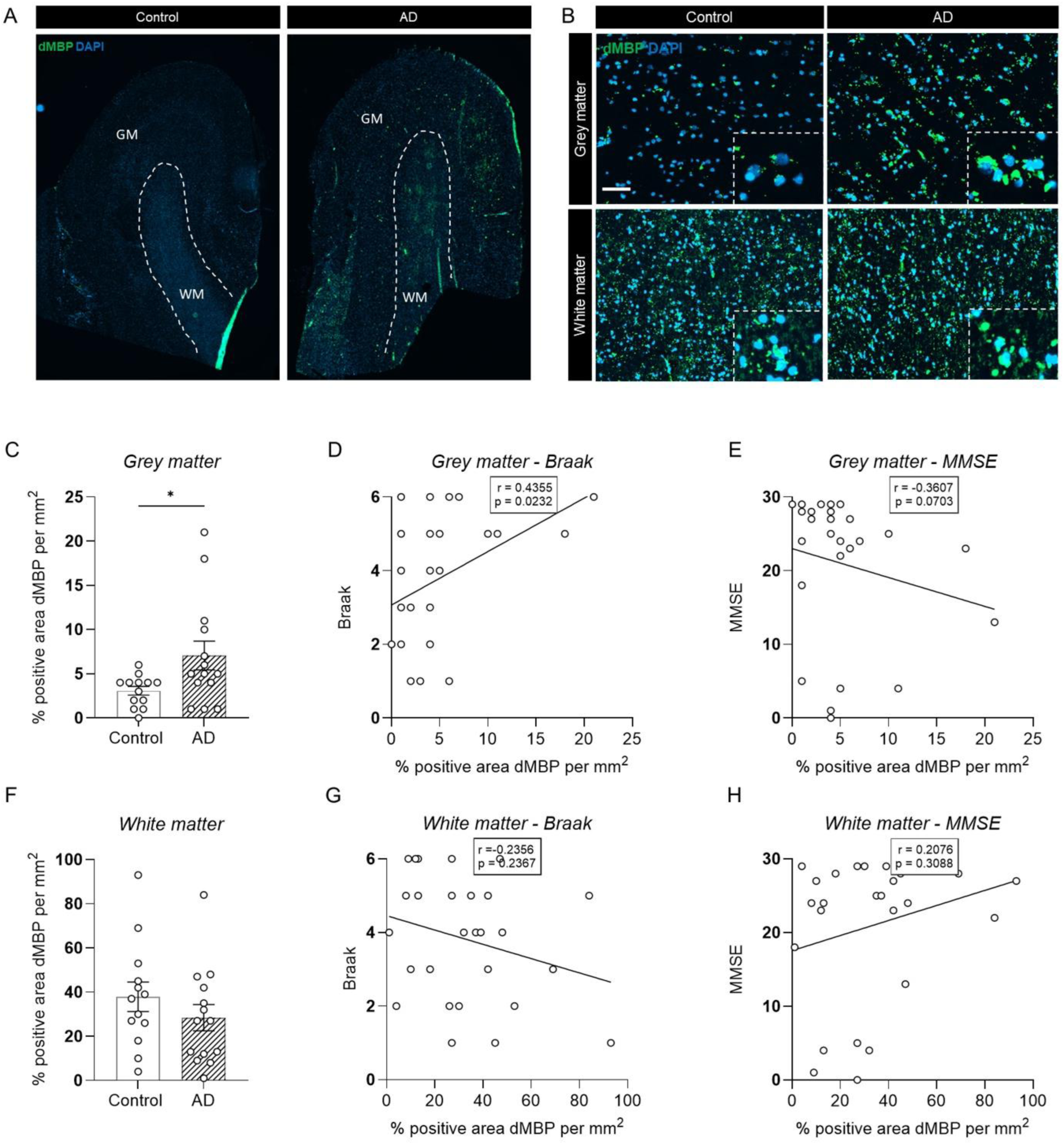
Increased myelin damage within AD grey matter. **(A–B)** Representative degraded MBP (dMBP) staining in brain sections from AD patients and age-matched controls, highlighting grey and white matter regions (scale bar: 50 µm). **(C, F)** Quantification of dMBP-positive area (fluorescence area per mm²) revealed a significant increase of myelin damage within the grey matter of AD patients compared to controls, while no difference was observed within white matter (n = 13–14, Mann–Whitney test, *p < 0.05). **(D–E)** Correlation analyses showed a significant correlation between grey matter dMBP area (%) and Braak stage and a trend towards a correlation between grey matter myelin damage and mini mental state examination (MMSE) score (Spearman correlation). **(G–H)** No significant correlations were observed between white matter dMBP area (%) and Braak stage or MMSE score (Spearman correlation).

### Loss of myelinating processes and mature oligodendrocytes in AD grey matter

Given the observed myelin loss and increased myelin damage in AD grey matter, we next investigated whether these changes were accompanied by alterations in the oligodendrocyte population. As an initial approach, we performed immunofluorescence staining for CNPase, an enzyme associated with myelin membranes that labels oligodendrocyte processes and myelin sheaths (Figure 3A,B). This enabled quantification of the area occupied by myelinating oligodendrocyte processes. We observed a reduction in CNPase immunoreactivity within both the grey and white matter of AD patients. Notably, only the decrease within the grey matter was associated with Braak stage (*P* = 0.0305, *r* = -0.4617) (Figure 3C–H).

**Figure 3.**
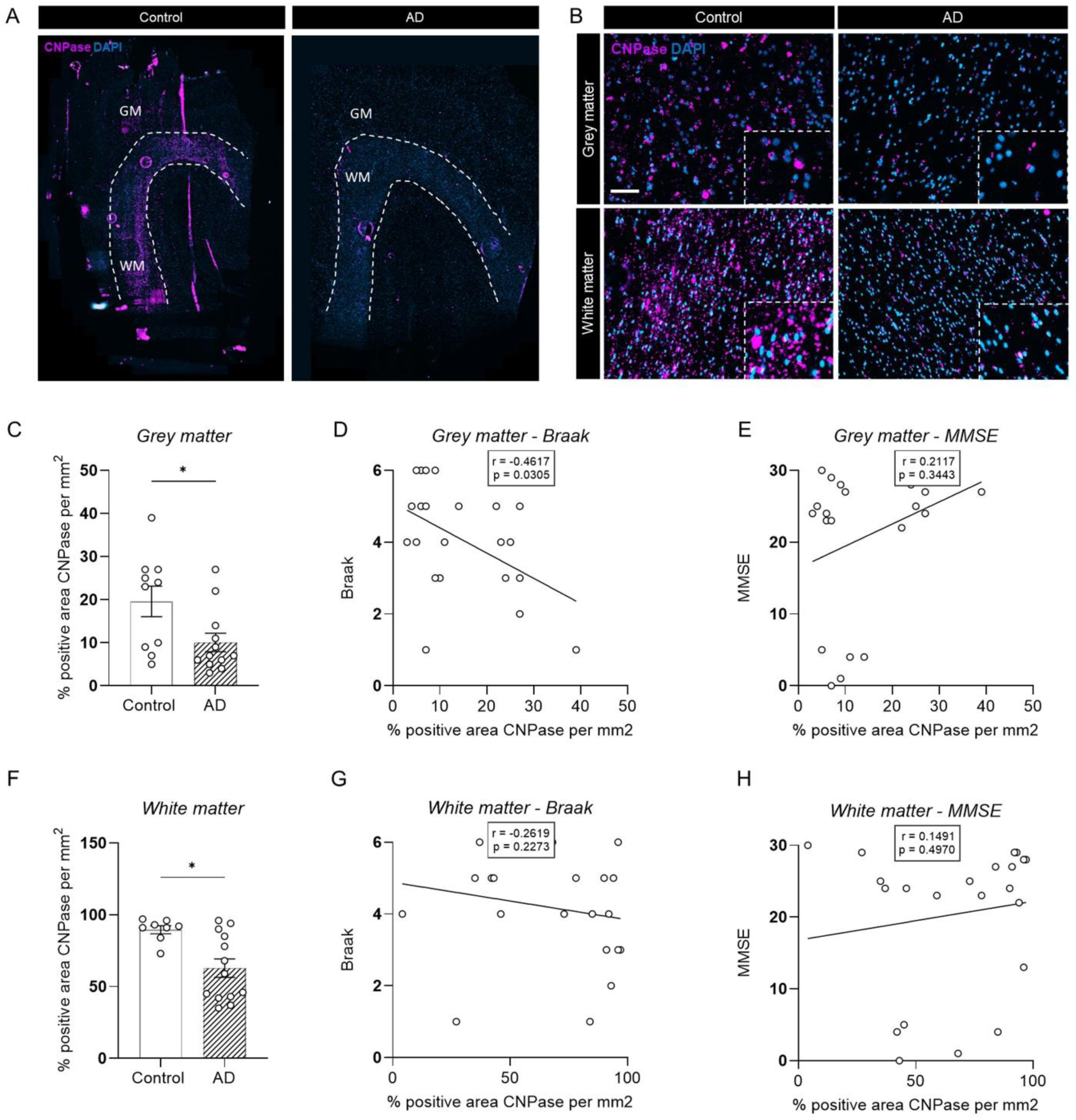
Reduced myelinating oligodendrocyte processes in AD are observed within both the grey and white matter, but only associate with disease stage within the grey matter. **(A–B)** Representative 2’,3’-cyclic nucleotide 3’-phosphodiesterase (CNPase) staining in brain sections from AD patients and age-matched controls, highlighting grey and white matter regions (scale bar: 50 µm). **(C, F)** Quantification of CNPase-positive area (fluorescence area per mm²) revealed a significant decrease within both grey and white matter of AD patients compared to controls (n = 10–12, Mann–Whitney test, *p < 0.05). **(D–E)** Correlation analyses revealed a significant association between grey matter CNPase-positive area (%) and Braak stage, but no correlation with mini mental state examination (MMSE) score (Spearman correlation). **(G–H)** In white matter, no significant correlations were observed between CNPase-positive area (%) and either Braak stage or MMSE score (Spearman correlation).

To specifically assess changes in mature oligodendrocytes, we performed co-immunostaining for CC1 and OLIG2 (Figure 4A,B). While the total number of oligodendrocyte lineage cells remained unchanged, quantification of CC1⁺/OLIG2⁺ cells revealed a significant reduction in mature oligodendrocytes within AD grey matter (*P* = 0.0216) (Figure 4C,D). This reduction did not correlate with Braak stage but was associated with MMSE scores (*P* = 0.0459, *r* = 0.4401) (Figure 4E,F). In contrast, no differences between AD and control samples were detected within white matter regions (Figure 4G–J).

**Figure 4.**
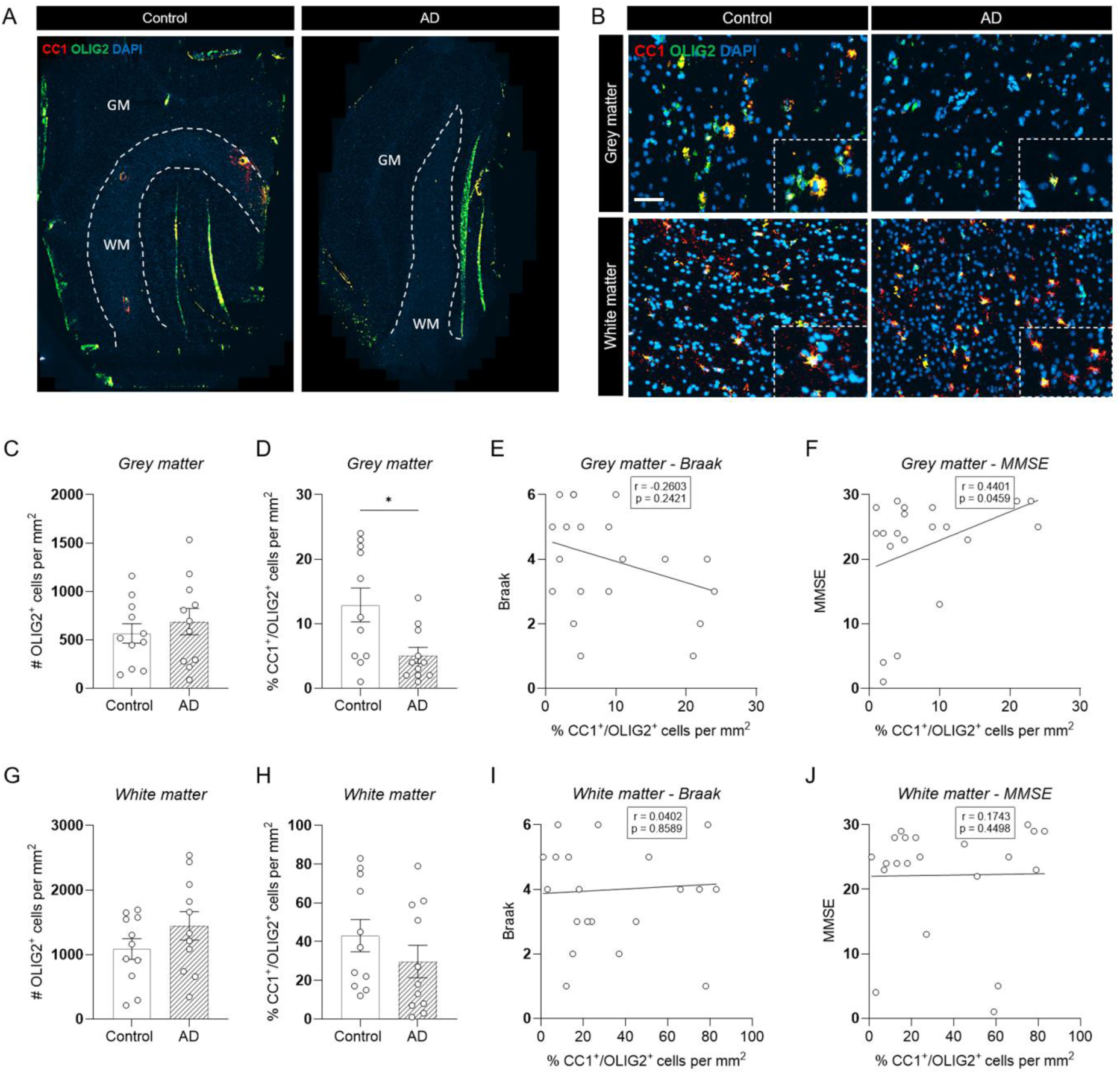
Loss of mature, myelinating oligodendrocytes within AD grey matter regions is associated with cognitive decline. **(A–B)** Representative Adenomatous Polyposis Coli (CC1)/Oligodendrocyte Transcription Factor 2 (OLIG2) staining in brain sections from AD patients and age-matched controls, highlighting grey and white matter regions (scale bar: 50 µm). **(C–D)** While the total number of OLIG2⁺ oligodendrocyte lineage cells remained unchanged, quantification of CC1⁺/OLIG2⁺ cells showed a significant reduction in AD grey matter compared to controls (n = 11; Mann–Whitney test, *p < 0.05). **(E–F)** Correlation analyses demonstrated a significant association between the number of CC1⁺/OLIG2⁺ cells in AD grey matter and mini mental state examination (MMSE) scores (Spearman correlation). **(G–J)** No differences were observed in white matter regions between AD and controls, and no correlations with Braak stage or MMSE scores were detected (Spearman correlation).

### Increased oligodendrogenesis and transcriptomic analysis hints towards insufficient compensatory myelin maintenance in AD grey matter

In multiple sclerosis (MS), chronic demyelination is often associated with impaired oligodendrogenesis, resulting in a failure of remyelination (15). To determine whether similar mechanisms are present in AD, we assessed the presence of newly formed oligodendrocytes using BCAS1 immunostaining (Figure 5A,B). Interestingly, we observed an increased proportion of BCAS1⁺ cells specifically within AD grey matter, suggesting an active, albeit potentially insufficient, attempt at myelin repair in these regions (Figure 5C). Notably, the abundance of BCAS1⁺ cells positively correlated with both Braak stage (*P* = 0.0069, *r* = 0.4987) (Figure 5D) and tangle load (*P* = 0.0116, *r* = 0.4702) (Supplementary Figure S2), paralleling the associations observed for MBP loss and dMBP accumulation. In contrast, no differences were detected between AD and control samples within white matter regions (Figure 5F–H). Given the significant correlation between BCAS1^+^ cell numbers, Braak stage, and tangle load, we next examined their relationship with tau pathology. To this end, we performed co-staining of AD samples for BCAS1 and AT8 (a marker of phosphorylated tau) and quantified the number of positive cells within concentric zones surrounding AT8^+^ regions (Figure 5I). We found that BCAS1^+^ cells were most abundant in the immediate peri-tau region, with their frequency progressively decreasing with increasing distance from AT8+ pathology (Figure 5J).

**Figure 5.**
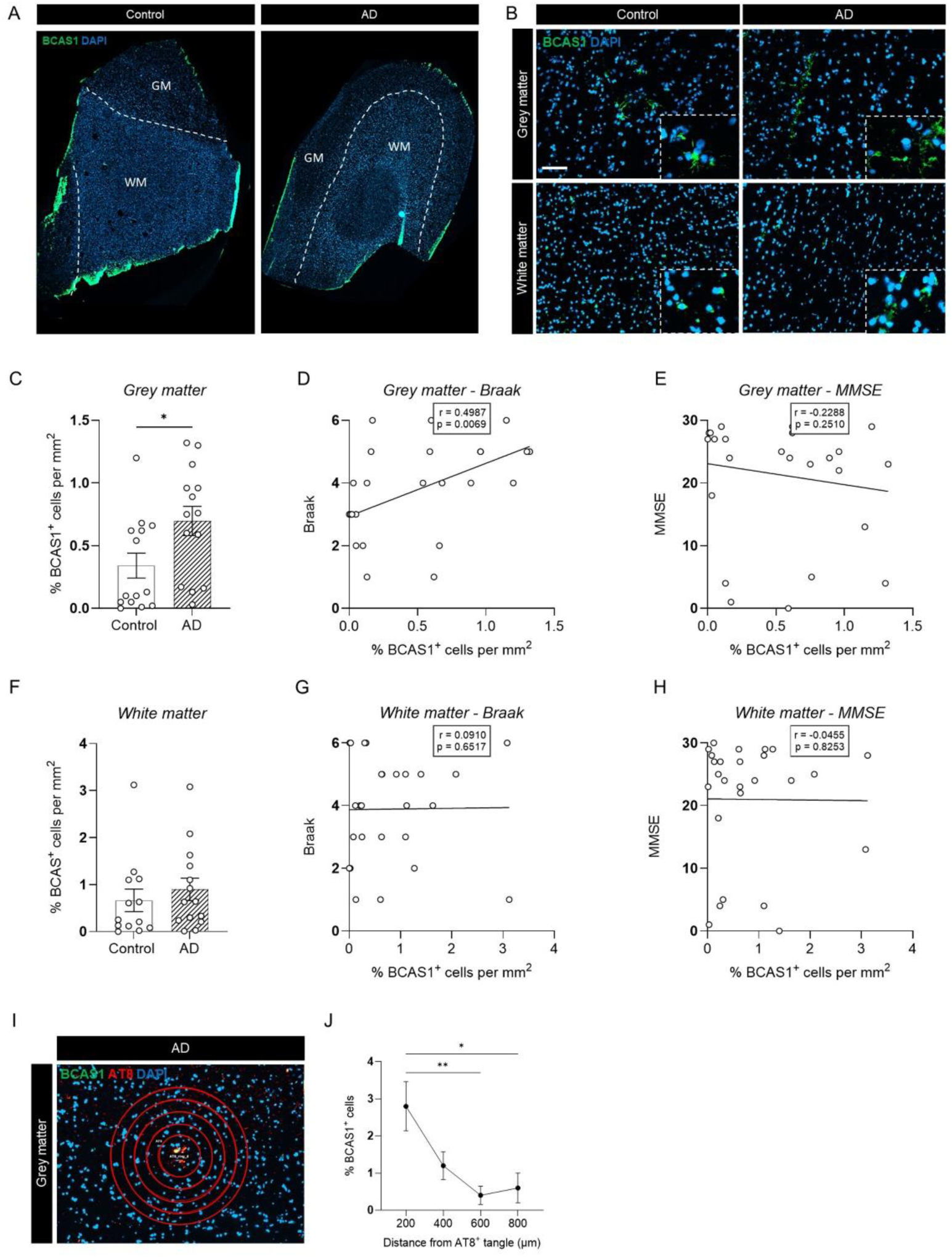
Increased oligodendrogenesis within AD grey matter regions is strongly associated with disease stage. **(A–B)** Representative Breast Carcinoma Amplified Sequence 1 (BCAS1) staining in brain sections from AD patients and age-matched controls, highlighting grey and white matter regions (scale bar: 50 µm). **(C, F)** Quantification of BCAS1⁺ cells (BCAS1⁺/DAPI per mm²) showed a significant increase within AD grey matter, with no difference observed within white matter (n = 14; Mann–Whitney test, *p < 0.05). **(D–E)** Correlation analyses demonstrated a significant association between the number of BCAS1⁺ cells within grey matter and Braak stage, with no correlation to mini mental state examination (MMSE) score (Spearman correlation). **(G–H)** In white matter, no significant correlations were observed between BCAS1⁺ cell abundance and either Braak stage or MMSE score (Spearman correlation). **(I)** Representative images illustrating spatial analysis of BCAS1⁺ cells relative to AT8⁺ tau pathology. Four concentric zones at 200 µm intervals were defined, and the percentage of BCAS1⁺ cells was quantified within each zone. **(J)** Quantitative analysis revealed increased BCAS1⁺ cell density in the immediate peri-tau microenvironment (n = 5, ordinary one-way ANOVA with Tukey’s multiple comparisons test, *p < 0.05, **p < 0.01).

In parallel, we performed bulk RNA sequencing on grey and white matter from control and AD brain samples. To further investigate oligodendrocyte-related transcriptional changes, we first generated a grey matter mature oligodendrocyte-specific gene set using a publicly available RNA sequencing dataset (GSE281006). We subsequently explored this signature within our bulk RNA sequencing dataset using ssGSEA. This analysis revealed significant enrichment of the grey matter mature oligodendrocyte signature in AD grey matter compared to controls (Figure 6A), whereas no enrichment was observed within AD white matter (Figure 6E).

**Figure 6.**
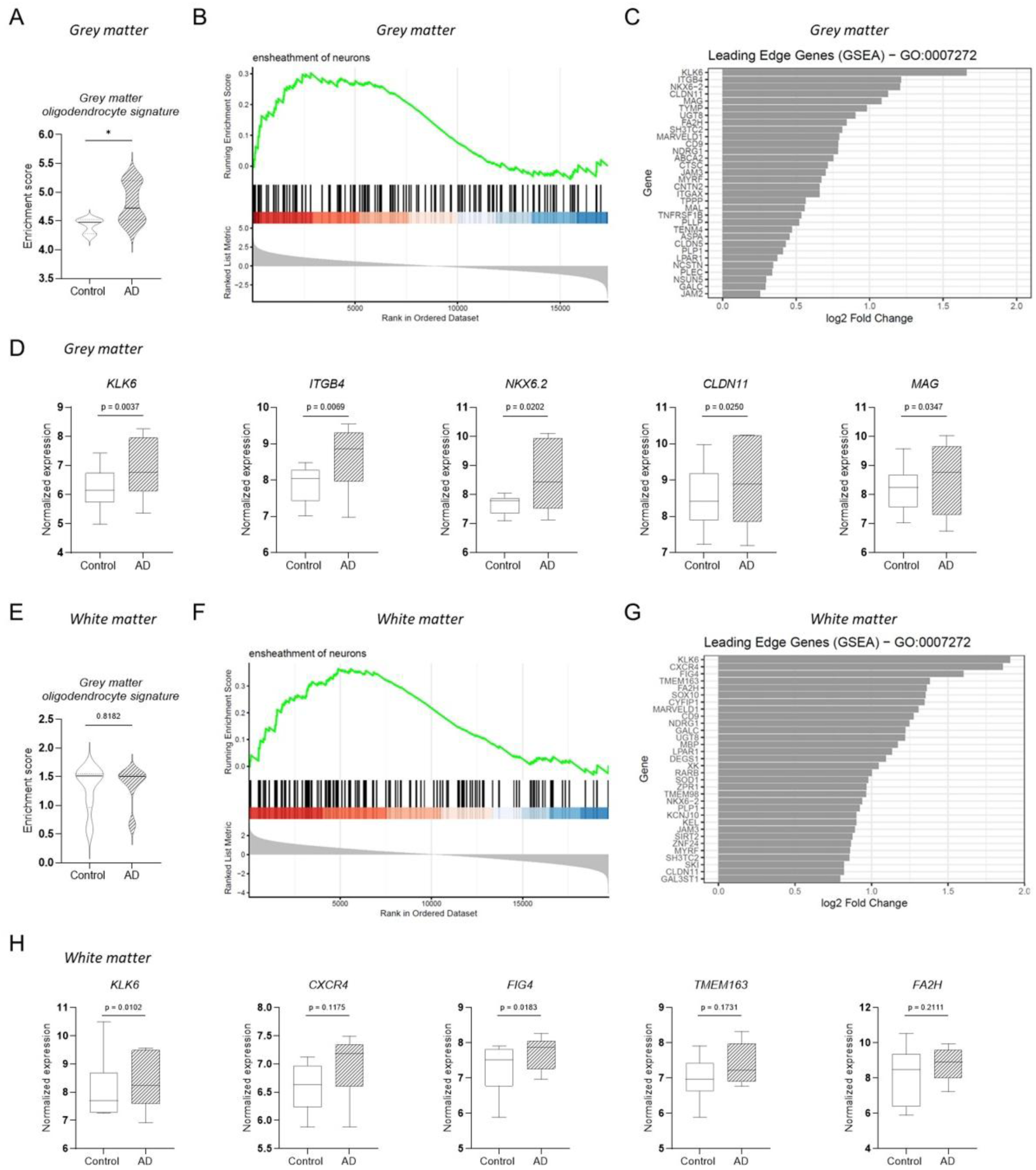
A myelin maintenance–remodeling transcriptional program within AD grey matter. **(A)** Single-sample gene set enrichment analysis (ssGSEA) of mature oligodendrocyte expression profiles derived from grey matter revealed significantly higher enrichment in AD grey matter samples compared to controls. **(B)** Gene set enrichment analysis (GSEA) of bulk transcriptomic data demonstrated significant enrichment of the GO term GO:0007272 (“ensheathment of neurons”) in AD grey matter relative to controls. **(C–D)** Leading-edge analysis identified coordinated upregulation of myelin-associated structural, adhesion, and remodeling genes, including *CLDN11, ITGB4, KLK6, MAG,* and *NKX6.2*, in AD grey matter. **(E)** In contrast to grey matter, white matter AD samples showed no enrichment of the grey matter mature oligodendrocyte expression profile. **(F)** Nevertheless, GSEA of white matter bulk transcriptomic data also revealed significant enrichment of the GO term GO:0007272 (“ensheathment of neurons”) in AD samples compared to controls. **(G–H)** Expression analysis in white matter further showed increased levels of genes associated with myelin turnover and cellular homeostasis, including *KLK6, CXCR4, FIG4, TMEM163,* and *FA2H*.

Following differential expression analysis of the bulk RNA sequencing data, we conducted GSEA, after which we focused specifically on oligodendrocyte- and myelination-related pathways. In particular, we examined the gene ontology term GO:0007272 (“ensheathment of neurons”), comprising 136 genes. This gene set was significantly enriched in both grey and white matter of AD patients compared to controls (Figure 6B,F).

Analysis of the leading-edge genes within this gene set revealed coordinated upregulation of myelin-associated structural, adhesion, and remodeling genes, with the most prominent changes observed in AD grey matter (Figure 6C–D). In grey matter, the top leading-edge genes included *CLDN11, ITGB4, MAG, NKX6.2*, and the proteolytic enzyme *KLK6*, consistent with a reactive oligodendrocyte transcriptional state associated with myelin instability and increased turnover. These findings suggest compensatory but inefficient myelin maintenance rather than effective remyelination. In white matter, increased expression of genes associated with myelin turnover and cellular homeostasis, including *KLK6 and FIG4,* further supported ongoing myelin remodeling processes in AD (Figure 6G–H). Notably, the leading-edge genes revealed higher fold changes in AD white matter, whereas greater statistical significance was observed in AD grey matter. Collectively, these findings suggest compensatory but inefficient myelin maintenance and remyelination in AD.

## Discussion

This study presents a comprehensive characterization of myelin alterations in post-mortem brain tissue from individuals with AD, discriminating between grey and white matter. We find that AD-associated loss of myelin and oligodendrocytes is predominantly localized within grey matter regions. These changes correlate with disease severity, as reflected by Braak stage, and with cognitive decline. In parallel, we detect signs of increased oligodendrogenesis and attempted myelin repair; however, these responses appear insufficient to compensate for the extent of ongoing degeneration.

This study is based on a unique post-mortem cohort comprising MTG tissue from AD patients and age-, sex-, and *APOE*-matched controls. The MTG is a region that is prominently affected in AD and has previously been shown, using magnetic resonance imaging (MRI), to exhibit demyelination (16, 17). Notably, the degree of myelin loss in this region has been associated with impairments in visuospatial cognitive function, underscoring its relevance to disease-related cognitive decline (17).

Importantly, our findings indicate that myelin degeneration in the MTG region of AD patients is predominantly confined to the grey matter. While white matter also shows signs of myelin alterations, similar changes are present in age-matched controls, suggesting that white matter myelin changes do not reflect AD-specific pathology. In contrast, grey matter myelin loss emerges as a more disease-specific feature. This selective vulnerability of grey matter oligodendrocytes has been suggested previously, where e.g. Tse *et al.* reported a loss of MYRF^+^ oligodendrocytes in grey, but not in white matter, in AD, and further linked this depletion to increased DNA damage (18). In addition, studies in surgically resected human brain tissue indicate that grey matter oligodendrocytes are intrinsically more susceptible to injury and exhibit transcriptomic signatures enriched for immune-related pathways and responses to environmental stressors, supporting the idea of an inherently distinct and more reactive oligodendrocyte population in grey matter (14).

In line with these observations, we show that grey matter myelin damage correlates with Braak stage and tau tangle burden, directly linking oligodendrocyte and myelin pathology to established neuropathological hallmarks of AD. This heightened vulnerability of AD grey matter may reflect its unique microenvironment, characterized by dense neuronal connectivity and a higher burden of toxic protein aggregation, which may increase metabolic and inflammatory stress on local myelinating cells and thereby exacerbate dysfunction and degeneration (19, 20). In this context, grey matter oligodendrocytes may represent a more fragile oligodendrocyte population compared to their white matter counterparts. Importantly, the extent of myelin disruption in the grey matter also shows a trend towards a correlation with cognitive impairment, as reflected in MMSE scores, supporting the idea that myelin integrity contributes to cognitive resilience (21, 22). Beyond its structural role, myelin in grey matter is increasingly recognized as a dynamic regulator of neuronal plasticity and circuit function through processes of adaptive myelination (23–25). Disruption of these processes may therefore impair neuronal network synchronization and cognitive adaptability, potentially contributing directly to functional decline in AD. This aligns with a model in which myelin alterations are not merely secondary to neurodegeneration but may actively influence disease progression and cognitive decline (26, 27).

In addition to the overall myelin damage, we observed a reduction in the number of mature, myelinating oligodendrocytes specifically within grey matter regions of AD brains, while the total pool of oligodendrocyte lineage cells remains largely preserved. This pattern suggests that AD preferentially disrupts oligodendrocyte maturation and/or survival, rather than causing a uniform depletion across the lineage (28). The maintenance of precursor populations may reflect a compensatory response, pointing to impaired lineage progression instead of global cell loss. Consistent with this interpretation, we detect signs of an attempted regenerative response, including an increase in BCAS1^+^ cells and transcriptomic signatures indicative of ongoing oligodendrogenesis and myelin repair (29, 30). BCAS1 is expressed by newly generated oligodendrocytes undergoing active myelination and is considered a marker of a transient intermediate stage during remyelination (31). The accumulation of BCAS1+ cells in the absence of a corresponding increase in mature myelinating oligodendrocytes may therefore suggest that repair is initiated but fails to progress to completion, either because differentiating oligodendrocytes become stalled in a transitional state or because they do not survive long enough to establish stable myelin sheaths (32). Consequently, despite evidence of endogenous repair attempts, this response appears insufficient to restore myelin integrity, indicating that regenerative mechanisms are either overwhelmed by continuous damage, or that (re)myelination within the diseased grey matter environment is intrinsically limited (14, 30, 33). Such a pattern differs from chronic demyelinating disorders such as multiple sclerosis, where remyelination attempts are more sustained, although ultimately unsuccessful (34, 35). We also observed a higher proportion of BCAS1+ cells in the immediate vicinity of phosphorylated tau aggregates than in more distal regions. Together with the consistent correlation between BCAS1+ cell abundance, tangle burden, and Braak stage, these findings suggest a potential mechanistic link between myelin vulnerability and tau aggregation.

To validate our findings at the molecular level, we performed bulk transcriptomic profiling of grey and white matter samples from both control and AD donors. Because bulk RNA-seq can be influenced by differences in cellular composition, we additionally leveraged an independent dataset of mature oligodendrocytes isolated from grey and white matter of healthy individuals (14). From this dataset, we derived a gene set based on differential expression between grey and white matter oligodendrocytes, which we then used to perform ssGSEA in our cohort. We observed a significant enrichment of this gene set in grey matter AD samples compared to matched controls, suggesting a disease-associated shift in oligodendrocyte state or reactivity within cortical regions. This was further supported by our GSEA results, which showed concordant changes. Although white matter exhibited larger fold changes in leading-edge genes, likely reflecting its higher density of myelinated fibers and oligodendrocytes, the grey matter showed a more coordinated transcriptional response. Together, these findings suggest that cortical oligodendrocyte dysfunction is a consistent feature of AD pathology, despite lower baseline myelin content in grey matter. Notably, the grey matter leading-edge genes showed a higher representation of structural and adhesion-related myelin components, including *CLDN11*, *MAG*, and *ITGB4*, as well as *KLK6*, a protease implicated in myelin remodeling and degeneration. Rather than reflecting effective remyelination, this transcriptional signature is more consistent with a reactive oligodendrocyte state characterized by increased myelin turnover and compensatory responses to ongoing damage. Such a state has been increasingly described in neurodegenerative disorders, where oligodendrocytes might upregulate myelin-associated programs but fail to fully restore myelin integrity or sustain axonal support (36–39). In this context, increased expression of ensheathment-related genes may reflect an attempt to stabilize vulnerable neuronal networks under chronic inflammatory, metabolic, and proteostatic stress.

Together, our findings support a model in which grey matter oligodendrocytes occupy a unique position of increased vulnerability in AD, contributing to region-specific myelin breakdown that is only partially compensated by endogenous repair mechanisms. Importantly, the association of myelin disruption with Braak stage, tau pathology, and cognitive decline further underscores the clinical relevance of oligodendrocyte and myelin pathology in AD progression. Our observations position oligodendrocyte dysfunction and maladaptive myelin remodeling as integral components of AD pathology and highlight grey matter myelin dynamics as a potential target for therapeutic intervention.

## Acknowledgements

This work was supported by Alzheimer Nederland (WE.03-2023-07), ZonMw Dementia Fellowship (10510022310003), and the ZonMw Dementia program (10510032120006).

## Author contributions

AT, DvdH and TV designed the study. AT took the lead in performing the experiments, analyzing and interpretation of the data, with critical input of EW, LK, MS, DvdH and TV. EW assisted in performing the experiments. AT, DvdH and TV drafted the manuscript. All the authors revised the manuscript and agreed to the final version of the manuscript.

## Supplementary Figures

**Supplementary Figure S1:**
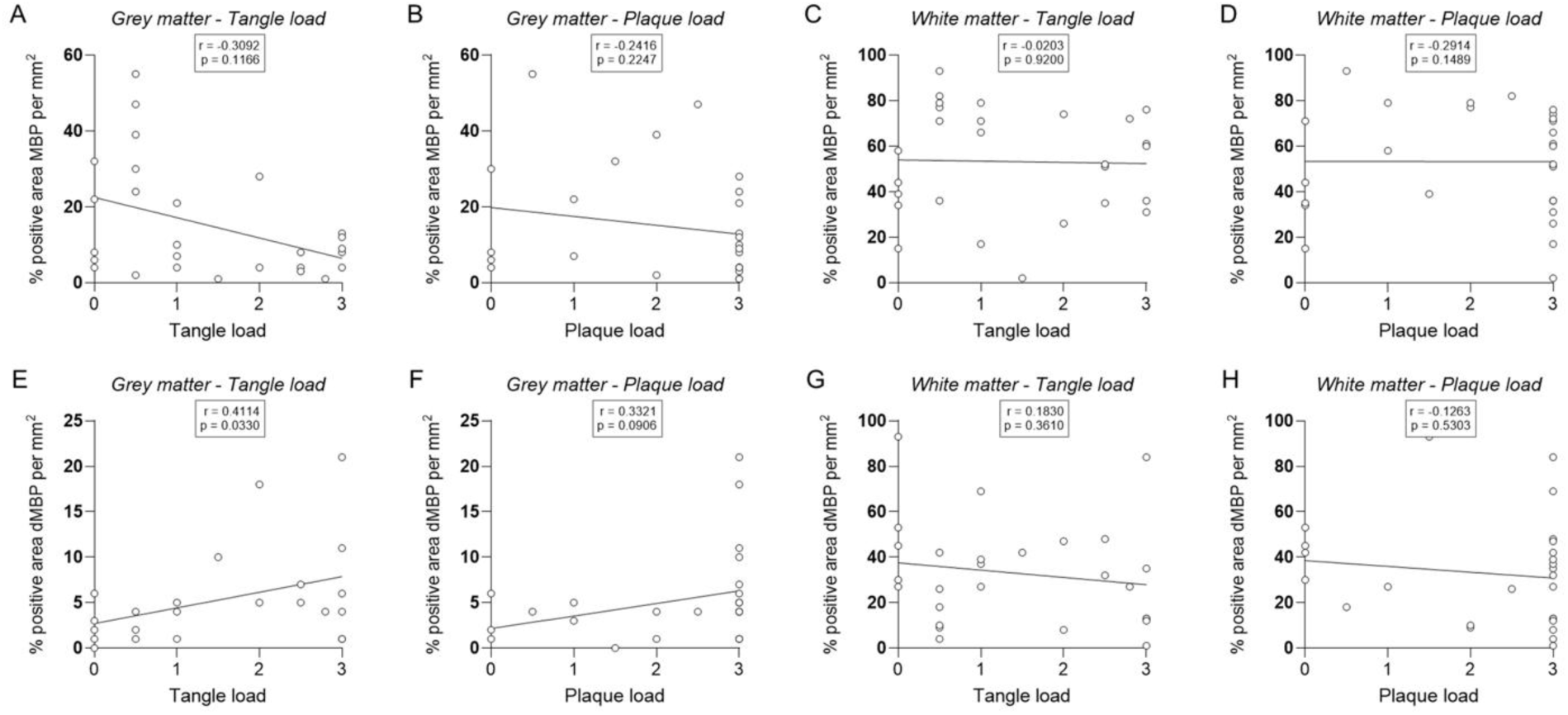
Spearman correlation analyses between MBP and dMBP levels and tangle and plaque burden in donor samples from grey matter (A, B, E, F) and white matter (C, D, G, H) regions.

**Supplementary Figure S2:**
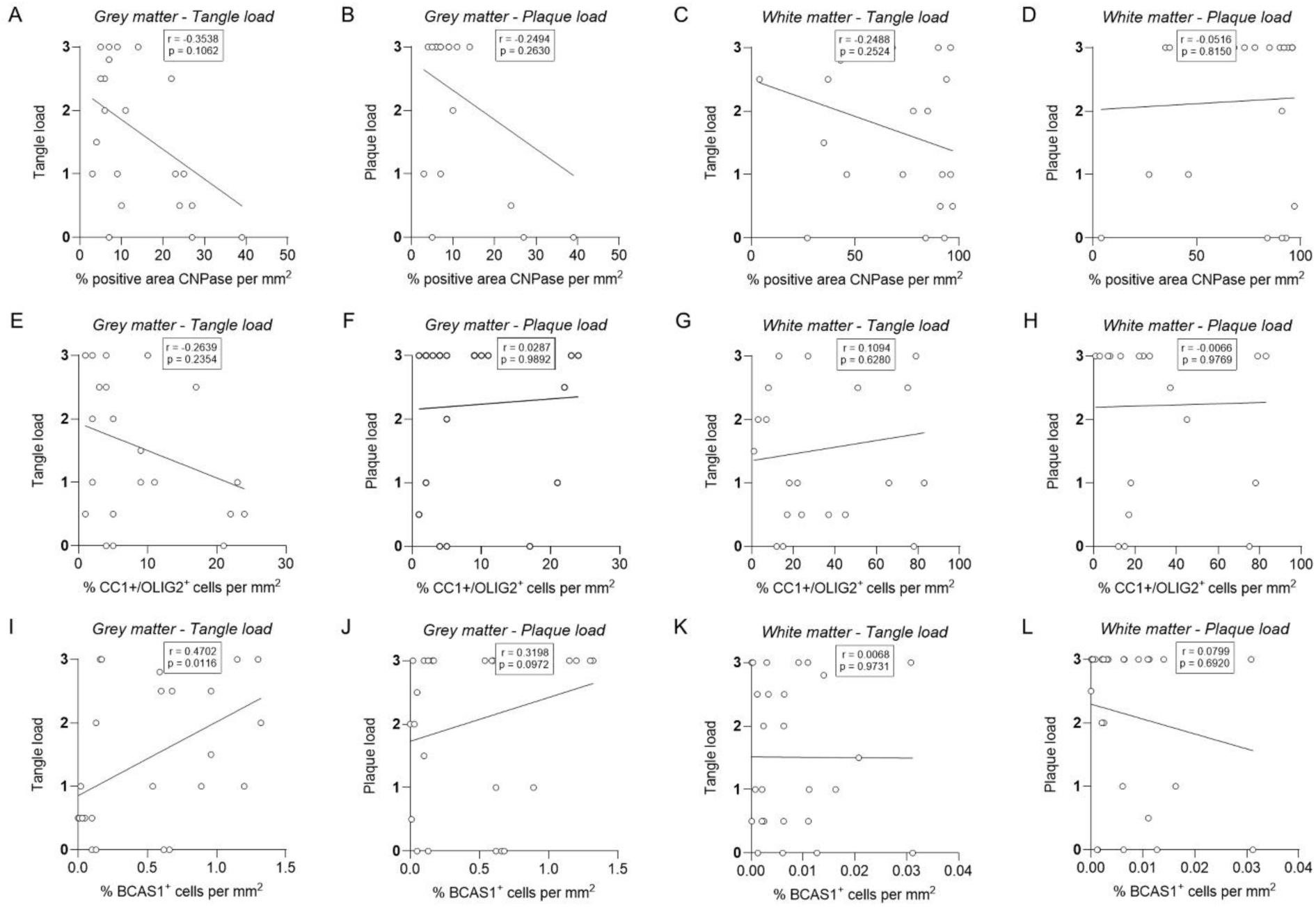
Spearman correlation analyses between CNPase, CC1/OLIG2, and BCAS1 levels and tangle and plaque burden in donor samples from grey matter (A, B, E, F, I, J) and white matter (C, D, G, H, K, L) regions.

## Supplementary Tables

**Supplementary Table S1:** Antibody list.

| Antigen | Antibody | Dilution |
| --- | --- | --- |
| MBP | MAB368 (Millipore) | 1:500 |
| dMBP | AB5864 (Millipore) | 1:2000 |
| CNPase | MAB326 (Millipore) | 1:1000 |
| CC1 | OP80 (Calbiochem) | 1:50 |
| OLIG2 | AF2418 (R&D Systems) | 1:50 |
| BCAS1 | 445 003 (SySy Antibodies) | 1:500 |
| AT8 | MN1020 (Thermofisher) | 1:1000 |

